# Membrane-bound cargo carried by teams of motors with heterogeneous velocities go faster and further than rigid cargo

**DOI:** 10.1101/2025.06.17.660035

**Authors:** Niranjan Sarpangala, Ajay Gopinathan

## Abstract

Intracellular transport by teams of molecular motors is an essential cell-biological process that ensures the proper distribution of organelles, and other materials within cells. These teams of motors cooperate and compete in complex ways to achieve desired transport velocity and runlength. *In-vitro* experiments have observed that coupling motors through a lipid membrane that mimics *in vivo* membrane-bound cargoes leads to a higher cargo velocity. However, the mechanisms behind this increase in lipid cargo velocity are unclear. Here we seek to understand these mechanisms using Brownian dynamics simulations. We show that an underlying heterogeneity in single motor velocity is essential for the increased velocity of lipid cargoes. Our simulations also show that while the runlengths of both rigid and lipid cargoes increase, and the velocities decrease, with an increase in the fraction of slower motors, lipid cargoes can travel faster and substantially further with the same degree of heterogeneity, suggesting functional advantages of motor velocity heterogeneity. Together, our work explains mechanisms behind previous experimental observations and generates new experimentally testable predictions on velocities and runlengths relevant for *in vivo* transport.

## Introduction

The spatial organization and correct functioning of eukaryotic cells require the active transport of materials and organelles from one part to another. This involves transport by teams of molecular motors along microtubule and actin filaments (12, 28) which depends on many physical characteristics, including cooperativity and competition between multiple motors(2, 3, 15–18) and interactions with the cellular environment and the morphology of the network (1, 9, 13, 19, 34). In addition, in cells, these teams of motors are coupled through the lipid bilayer that typically encloses the cargo. This lipid bilayer is present in most cargoes like lipid droplets, peroxisomes, synaptic vesicles, and organelles such as mitochondria. Recent works have shown that such coupling through the lipid membrane, and the the resulting diffusion of motors on the cargo surface influence multimotor functioning (4– 6, 8, 14, 21–24, 26, 31, 33, 38) in various ways. In particular, studies have shown that cargo velocity (22, 26), runlength (average distance traveled) and mean binding rate (14, 23, 33) can change due to the presence of the membrane, as membrane diffusion helps to reduce mechanical interference (33) and leads to dynamic motor clustering. Interestingly, microstructures in the membrane such as lipid rafts can lead to additional clustering of motors (31)

In addition to the complexities of motor-motor coupling through the membrane, the motor teams in cells are found to consist of motors of different types. For example, it is known that melanosomes in *Xenopus* melanophores are carried by kinesins, dyneins, and also myosin-V (20), while intraflagellar transport of IFT particles depends on kinesin-II and osm-3 kinesin (29, 30, 35). There are also abundant examples of the presence of both dyneins and kinesins on the same cargo, which leads to bidirectional transport on microtubules (8, 11). It is still unclear how cells can achieve desired transport when the individual motors have different transport properties, such as velocity, binding and unbinding rate, and even polarity.

This transport by non-identical motor teams has been studied experimentally and also theoretically. It is shown in the microtubule gliding assay experiments and models that the predicted behavior of teams of motors is sensitive to their single motor properties (3). However, the insights from the planar geometry in gliding assay systems cannot be directly translated into the spherical cargo geometry relevant for transport in cells and can lead to different expectations for run length and velocity (33). Hence, one has to complement these studies with studies on spherical cargo geometry for a comprehensive understanding of transport by heterogeneous teams of motors through complex cellular environments.

Experiments on spherical cargo geometry often focus on two quantities, the cargo runlength and cargo velocity since they are the two transport quantities of most relevance *in vivo*. For example, if we consider the transport of signaling molecules, then cargo velocity might be more important as it determines how rapidly signaling molecules can reach the target (25). In this context, it is interesting to note that cargoes that have a fluid surface have been observed to have a higher velocity than those that have rigid or gel-like surfaces when carried by teams of kinesins (22) as well as myosin-V(26) in *in vitro* experiments. These studies also propose two possible mechanisms that could lead to this speedup, namely,(i) the delay in strain generation between motors of different types and (ii) the enhanced detachment of lagging motors and recentering of cargo. However, it is unclear whether these mechanisms explain the observed speedup quantitatively and what the relative contribution of these two mechanisms to the overall speedup is. We wanted to find a minimal computational model that shows this velocity difference and thereby understand the mechanisms underlying higher lipid cargo velocity.

Our previous simulations of lipid cargo transport (33) did not show a statistically significant difference in rigid and lipid cargo velocities. We reasoned that this could be because the force-velocity dependence we assumed in our computational model may not be sensitive enough to lead to differences in cargo velocity with changes in cargo surface fluidity. It may also be due to the heterogeneity in single motor velocity often seen in mixtures of motors purified from cells (26, 32). In fact, it was suspected that this heterogeneity in single motor velocity could lead to the observed higher lipid cargo velocity (22), although it was not explained how heterogeneity could lead to higher velocity.

In this study, we performed cargo transport simulations by motor teams with varying levels of heterogeneity in their individual motor velocities. Our findings indicate that a substantial degree of heterogeneity in single motor velocity is crucial for achieving higher lipid cargo velocities. We elucidated this increased velocity through an analysis of cargo and motor dynamics, the velocity contribution from cargo recentering mechanisms, and the delay in strain generation and found that strain generation delay was the dominant mechanism. As expected, cargo velocity also decreases with an increasing fraction of slower motors. Furthermore, we investigated the impact of heterogeneity in motor velocity on runlength and demonstrated that lipid cargoes have a significantly higher runlength than rigid cargo, which increases with an increasing fraction of slower motors. This means that they can also traverse a specified distance with a smaller proportion of slower motors thereby maintaining a higher overall cargo velocity.

## Methods

### Brownian Dynamics Model

In this study, we modified the Brownian dynamics computational model of multi-kinesin transport used in our study (33), and incorporated heterogeneity in the single motor velocities. Briefly, we consider a spherical cargo with a fluid surface transported by multiple kinesin-1 motors. Each cargo had a fixed number of motors (*N*), which diffuse on the cargo surface. Each cargo is initialized with the *N* motors randomly placed on the spherical surface. In cells, the number of motors on a cargo need not be fixed, which was shown to influence the cargo run length (36). However, we choose to ignore this in our study because we are focused on the impact of the presence of the membrane on the cargo surface and motor velocity heterogeneity keeping all other variables fixed. These motors diffuse on the cargo surface with a diffusion constant, *D* which is a parameter in our model. For rigid cargo, we set *D*=0, and for fluid cargo, for simplicity, we used *D* = 1 *µm*^2^*s*^−1^ which is a typical diffusion constant of motors on membranes (10). Kinesin motors that are attached to the microtubule were modeled as one-sided springs (18) with one end (head) on the microtubule and the other end (anchor point) on the cargo. The kinesins exert a spring-like force when extended beyond their rest length; when compressed below this length, the kinesins exert no force. We considered the force and ATP dependence of the bound motor’s off-rate and stepping rate. We also incorporated the rotational diffusion of the cargo due to the torque from motor forces and thermal fluctuations. The center of mass of the cargo was updated according to overdamped Langevin dynamics.

To take into account the heterogeneity in motor velocities, we considered experimentally observed heterogeneity in motor populations. Specifically, we used a normal distribution and a bi-delta distribution of velocities, as explained in the results section. We assigned to each motor a mean unloaded motor velocity *v*_*o*,*mot*_, which was drawn randomly from the chosen probability distribution. In our model, the microscopic unbinding rates of motors are inversely proportional to their stepping rates. Thus slower motors effectively spend longer times associated with microtubules. Please see supplemental information (Section I) for more details on the computational model.

## Results

### Heterogeneity is essential to explain experimentally observed differences in rigid and lipid cargo velocities

We wanted to first find a computational model with the least set of assumptions that shows a velocity difference between rigid and lipid cargoes. While previous models have shown this velocity difference, (26) they had a few assumptions. First, it was assumed that every step of a motor results in a spike in its force. Further, it was assumed that if the motor is attached to lipid cargo, the tangential component of this force relaxes quickly because of the sliding of motor attachment point on the membrane, and if it is a rigid (or gel-state) cargo then such force relaxation is much slower. While our previous computational model (33) with Langevin dynamics of motors on the cargo surface showed that there is indeed a reduction in forces experienced by motors in lipid cargo, it is only true statistically, and force trajectories don’t show a spike and eventual relaxation every time a motor steps. Second, it was assumed that the center of mass of the cargo fluctuates about a point that is the mean of front and rear-most motor positions, which need not necessarily be true. We were curious to know if we are able to see the velocity difference between rigid and lipid cargo in a 3-dimensional Brownian dynamics simulation of cargo transport by teams of motors independent of additional assumptions of motor force relaxation and cargo recentering.

Surprisingly, our initial simulations of 3-dimensional cargo transport by teams of motors, reported earlier (33) didn’t show any statistically significant difference between rigid and lipid cargoes. We reasoned that this independence of cargo velocity from membrane fluidity could be because the force-velocity curve we used in our simulation was not sensitive enough or, as previous works (22) pointed out, this may be due to the presence of heterogeneity in motor velocities in the team.

To test if heterogeneity could lead to differences in lipid and rigid cargo velocities, we first introduced heterogeneity by incorporating a normal distribution for the single motor velocity with a standard deviation of *σ* = 20 *nm s*^−1^ (37) (Fig. 2a). In this case as well, no statistically significant difference was seen between rigid and lipid cargo velocities (Fig. 2b). We thought this could be because the heterogeneity that we considered wasn’t sufficient to cause a significant difference in cargo velocities. In fact, previous experimental works have reported a higher relative heterogeneity in single motor velocity in teams of kinesins (32) and also in teams of myosins (26) as quantified by the spread in the single motor velocity distribution. When we used a high heterogeneity normal distribution with *σ* =250 *nm s*^−1^ (Fig. 2a), we observed a higher velocity for lipid cargoes (Fig. 2b). This shows that heterogeneity in single motor velocity could lead to a difference in rigid and lipid cargo velocity. To confirm that heterogeneity in single motor velocity leads to the velocity difference, we tried another case of heterogeneity, seen in kinesin-I motors purified from drosophila embryos (32). These motors showed two subpopulations with velocities of 350 *nm s*^−1^ and 800 *nm s*^−1^ at a population ratio of 0.3:0.7, respectively. We approximated this distribution by a bi-delta distribution (Fig. 2a). This case also resulted in a higher velocity of lipid cargoes, confirming that heterogeneity in single motor velocity leads to higher velocity of lipid cargoes (Fig. 2b).

**Figure 1.**
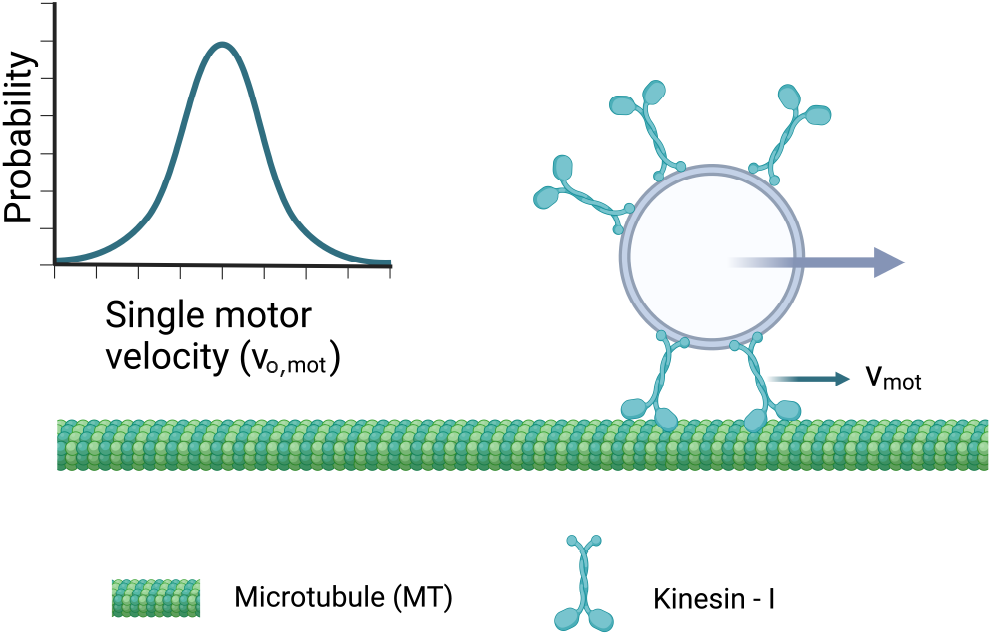
Schematic of the computational model. The velocity of a motor *v*_*mot*_ depends on the force that it is experiencing and its intrinsic unloaded motor velocity *v*_*o*,*mot*_ . The unloaded motor velocity *v*_*o*,*mot*_ is different for each motor, and they are drawn randomly from a given probability distribution like the one shown on the left. Schematics created with Biorender.com

**Figure 2.**
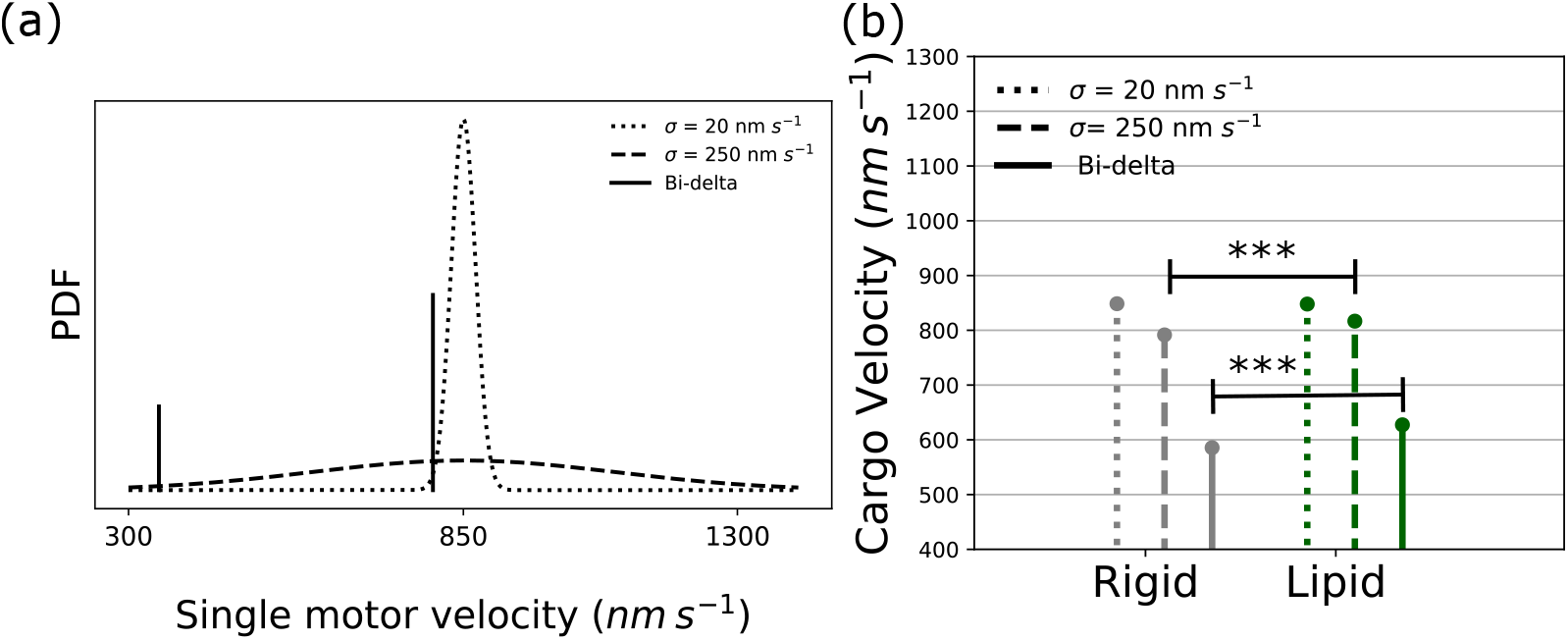
Heterogeneity is essential to explain experimentally observed differences in rigid and lipid cargo velocities. (a) The three different single motor velocity distributions considered in our study, normal distributions with *σ*= 20 nm/s, *σ* = 250 nm/s and bi-delta distribution - two different velocity populations with velocities 350 nm/s and 800 nm/s at the population ration of 0.3:0.7. When we initialize cargoes, we assign to each motor its unloaded velocity drawn from these distributions. (b) The resultant cargo velocity for three different input single motor velocity distribution. To measure velocity, we obtained cargo position data from the simulations of the transport of cargo by teams of motors with the given single motor velocity distribution. For each cargo run, we recorded the cargo position data at a sampling rate of 100 *s*^−1^. 200 such cargo runs were considered for each parameter set. Cargo velocity was then measured as the ratio of mean displacement along the x-axis in a given time window to *δt* (we took *δt* = 0.1 s). Error bars denote the standard error of the mean (error bars are very small and hence not visible in the figure). For all cases, the number of motors on the cargo was *N* = 16 and ATP concentration in the medium, [ATP] = 2 mM. For lipid cargoes, we considered a motor diffusion constant of *D* = 1 *µm*^2^*s*^−1^.

Our observations compare well with previous experiments on cargo transport by teams of kinesin-1 motors (22). Here it was found that the velocity difference between rigid and lipid cargo is an increasing function of the motility fraction (a proxy for the number of motors on the cargo). This velocity difference varies from about 25 *nm s*^−1^ at low motility fractions to about 150 *nm s*^−1^ at a very high motility fractions (22). Interestingly, the value of velocity difference that we observe, about 30 *nm s*^−1^ is within this range. In addition to that, we observe velocity difference between rigid and lipid only at high number of motors on the cargo (see supplemental information, Fig. S1) consistent with experimental observations. However, experimental data (22) shows the velocity difference between rigid and lipid arises not because the velocity of lipid cargo increases with increase in motility fraction but rather because the velocity of rigid cargo decreases while the lipid cargo velocity remains constant. Thus one could argue that the lipid cargo is not moving faster than average, but in fact rigid cargo is moving slower. This points to the possibility that other mechanisms might be at play, possibly the interference between motors (7, 33) to cause rigid cargoes to move slower than lipid cargoes in teams of kinesins.

However, experiments on teams of myosin Va (26) show that lipid cargoes move faster than average single motor velocity, indicating that the velocity difference between lipid and rigid (gel state) is due to the additional velocity that lipid cargoes gain due to the underlying motor dynamics. This is in agreement with our simulations. However, in this experiment, it was observed that the lipid cargo velocity decreases with an increase in the number of motors on the cargo. This might be because of the actin site exclusion and resulting jam between myosin Va motors. Our model doesn’t take into account the filament site exclusion - the fact that one site can be occupied by only one motor at a time. While neglecting site exclusion is reasonable for kinesin dynamics of microtubules that have multiple protofilaments, site exclusion might be important for actin filaments that have only two protofilaments. This might explain the discrepancy between our simulations and experimental observation of velocity at high motor densities. In this experiment (26), it was also noted that the velocity of lipid cargo increases with increasing cargo size, while the rigid cargo velocity remains constant. Although we see a trend that confirms this experimental observation (see supplemental information, Fig. S2), there are also high fluctuations that make it difficult to say with certainty whether the lipid cargo velocity increases with an increase in cargo size.

Overall, our simulations reproduce the experimentally observed increase in lipid cargo velocity and show that the underlying heterogeneity in single motor velocity is crucial for this phenomenon.

### Delay in strain generation between slow and fast motors due to the presence of membrane explains the observed increase in lipid cargo velocity

We next wanted to understand the mechanisms that lead to higher lipid cargo velocities in the presence of motor speed heterogeneity. For simplicity, we focused on the bi-delta case (mixture of slow, 350 *nm s*^−1^ and fast, 800 *nm s*^−1^ motors at a population ratio of 0.3:0.7). Since we had a simple case, a mixture of slow and fast motors with defined velocities, we hoped to find an analytical estimate of cargo velocities as a starting point for our explanations. If we assume that only one motor is bound at a time, this motor could be slow or fast; the probability of it being slow or fast depends on the population ratio and the mean lifetime of these motors. We then computed the average of slow and fast motor velocities weighted by the corresponding lifetime and population of slow and fast motors, which is *v*_*est*_ = 577.32 *nm s*^−1^ (see supplemental information, Section II for more details), which is lower than the observed cargo velocities of 598 ±1.4 *nm s*^−1^ (for Rigid cargo) and 633± 1.7 *nm s*^−1^ (for Lipid cargo). It could be because, most of the time, more than one motor is bound, and faster motors could be enhancing the detachment of slower motors. We looked at the probability distribution of the number of bound motors, and only 30 to 40 % of the time, it is singly bound (see supplemental information, Fig. S3). If we consider that more than one motor is bound, it becomes important to take the mechanics into account since the velocity difference between motors could induce a strain between them which could impact the motor and cargo velocity. In fact, forces experienced by slow and fast motors do show such a strain build-up between slow and fast motors; when the lagging slow motor detaches, this additional strain experienced by the fast motor drops (see supplemental information, Fig. S4).

In previous work, it has been emphasized that the increase in the cargo velocity of lipids is due to the preferential detachment of the lagging motors, and after a lagging motor detaches the cargo quickly and recenters, giving an additional boost to the cargo velocity (the mechanism is illustrated in Fig.3b). It was suggested that there is a higher contribution from such a cargo recentering effect for lipid cargo since the lipid membrane allows for relaxation of the tangential component of force, resulting in the situation where a motor that detaches preferentially has a force normal to the cargo surface (26).Since our simulation allowed us to analyze the dynamics at a single motor level, it was straightforward to analyze the cargo dynamics right after a motor detached from the microtubule and thereby test the hypothesis.

We measured the velocity of the cargo immediately after a motor unbinds, which is observed to decay exponentially as the time window size used for measuring the velocity increases (see supplemental information, Fig. S5). We fit this data to *v*_cargo_+Δ*x* _*f*_ /*t* to obtain the mean flopping distance Δ*x* _*f*_ ±*δx* _*f*_ (using scipy.optimize.curve_fit, *δx* is the one standard deviation error in Δ*x* _*f*_). *v*_cargo_ is the average cargo velocity. Surprisingly, the flopping distance for rigid cargo is higher than for lipid cargo. We also measured the number of flops, *i*.*e*., the number of motor detachments, *N* _*f*_, and the total cargo simulation time, *T* (equal to the cumulative lifetime of 200 cargo runs). Then we estimated the mean contribution to cargo velocity from the cargo recentering mechanism as Δ*v*_flop_ = *N* _*f*_ Δ*x* _*f*_ /*T* and the error as *δv*_flop_ = *N* _*f*_ *δx* _*f*_ /*T* . Interestingly, the contribution to the velocity from this cargo recentering was less for lipid cargo than for rigid cargo (Fig. 3(b)). Hence, the cargo recentering mechanism does not explain the observed velocity difference between rigid and lipid cargoes. We also added the contribution from cargo recentering (Δ*v* _*f lop*_) to our estimation of cargo velocity (*v*_*est*_), and obtained a value of about 593.59± 0.7 *nm*/ *s* for rigid cargo and 589.6 ± 0.3 *nm* /*s* for lipid cargo. This value is close to the observed value for rigid cargo, but not for lipid cargo.

**Figure 3.**
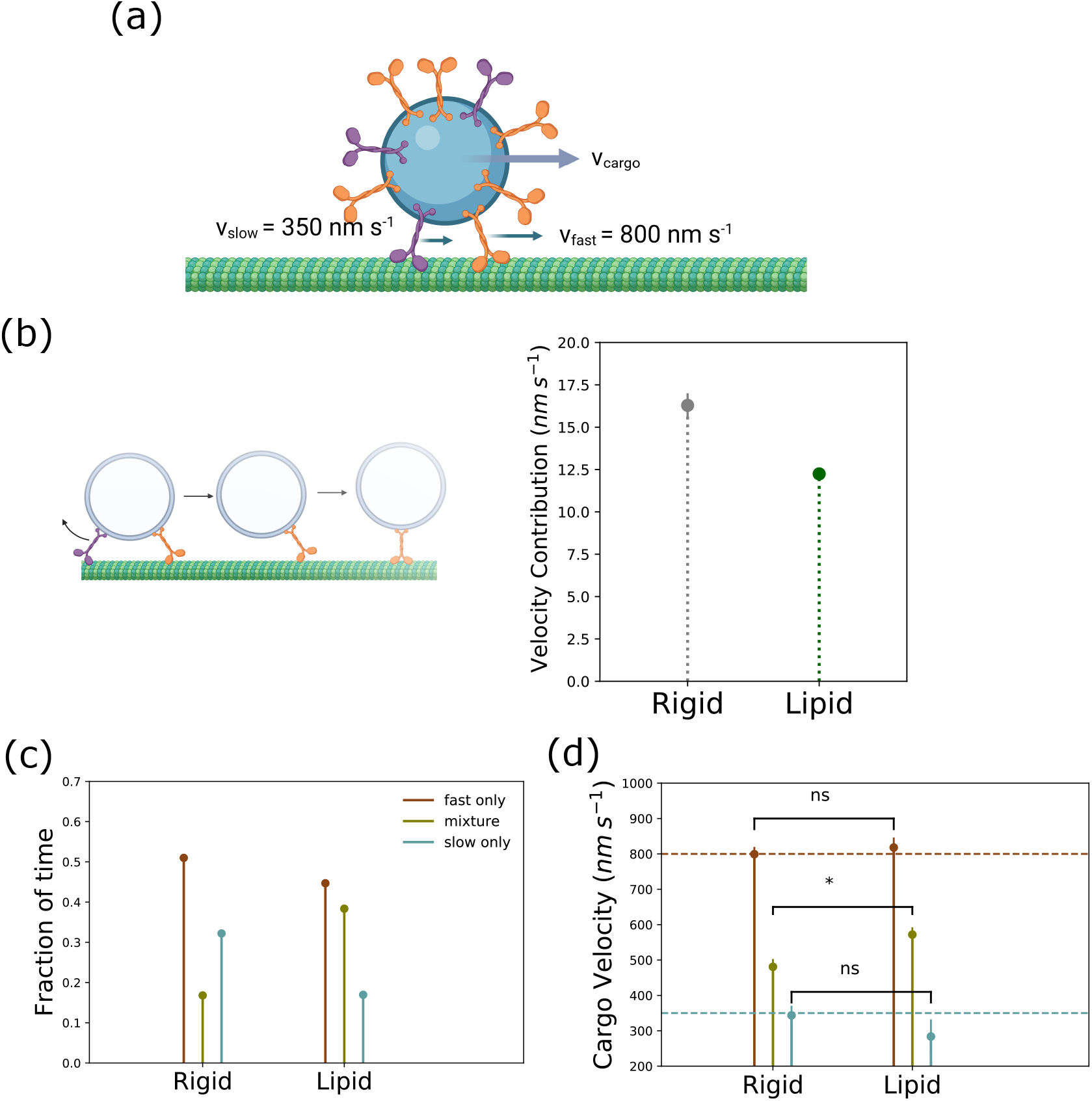
Delay in strain generation between slow and fast motors explains the observed increase in lipid cargo velocity. (a) Schematic of the cargo-motor system considered in this section. Single motor velocities of motors were 350 nm/s (slow) and 800 nm/s (fast) at a population ratio of 0.3 to 0.7 (a bi-delta distribution) (b) Illustration of [left] and quantification of the contribution to cargo velocities from [right] cargo recentering mechanism. Error bars are calculated from the error propagation method described in the text. (c & d) Evidence for the delay in strain generation between slow and fast motors in lipid cargoes;(c) Fraction of time and (d) Mean cargo velocity when cargo is being carried by only fast, only slow, and a mixture of slow and fast motors. To get data in (c) and (d), we first ran simulations of cargo transport with N=16, [ATP]= 2 *mM* and recorded data at a sampling rate of 100 *s*^−1^ for 200 independent cargo runs each for rigid and lipid cargo. From this data set, we filtered out time windows where microtubule-bound motors were (i) all fast motors, (ii) all slow motors, and (iii) a mixture of fast and slow motors. To get fraction of time data in (s), we computed ratio of the cumulative size for each filtered group to the total size of the unfiltered data. To get the velocity data in (d), we measured the cargo velocity by computing mean displacements in short time windows (Δ*t* =0.01 s) among each filtered group. Error bars indicate the standard error of the mean. p values were computed from the students-t test. * represents p *≤* 0.05 and ns represents p > 0.05. Cartoons in (a) and (b) created with Biorender.com.

To resolve this discrepancy, we made a closer analysis of cargo and motor dynamics when multiple motors are working together. Since the motors move at different speeds, we expect them to compete mechanically with each other. For teams of identical motors, it was shown that the fluidity of the membrane reduces mechanical competition (33). We expected that when we have teams of slow and fast motors, the lipid membrane could also reduce the mechanical competition, thereby allowing slow and fast motors to spend more time working together. In other words, the lipid membrane could delay strain generation (22).

In order to test this hypothesis, it is important to look at the typical composition of bound motors teams, whether it is predominantly fast motors, predominantly slow motors or a more even mixture of fast and slow mixtures. This is not only a function of the lifetime and population ratio of motor teams on the cargo but also depends on the mechanical interactions between motors. The fraction of the time cargo is carried by all fast motors, all slow, and the mixture that we measured from the simulated data is given in Fig. 3c. It can be seen that the fraction of time the cargo is carried by a mixture of the slow and fast motors is more in the case of lipid cargo compared to rigid cargo. This means that lipid membrane allows slow and fast motors to spend more time associated with the microtubule simultaneously. This is in agreement with our expectations of reduced mechanical interference in lipid cargoes.

Then we were curious to know what is the cargo velocity when being carried by slow only, fast only or a combination of some fast and some slow microtubule-bound motors. While it is easy to predict what the mean cargo velocity should be when all microtubule-bound motors are fast (or when all are slow motors), it is unclear what the cargo speed should be when some slow and some fast motors are simultaneously bound to the microtubule. One could argue that if any of the MT bound motors is a slow motor, then the entire cargo should be moving with slow motor velocity. However, analysis of simulation data indicates that the cargo velocities when some of the MT bound motors are slow, are higher than the individual slow motor velocity (Fig. 3d).

This higher velocity could be because these slow and fast microtubule-bound motors may not always be under tension. This is confirmed by the distribution of motor forces experienced by slow and fast motors (supplemental information, Fig. S6) which indicates that slow and fast motors experience low forces (i.e. any forces below 5.25 pN - for a hindering load of 5.25 pN, the fast motor velocity reduces to slow motor velocity) for a sufficient fraction of time during the time they associated with microtubules. We believe that as long as motors experience forces less than 5.25 pN, the fast motors move at velocities higher than the slow motor velocity. Consequently, the cargo center of mass also moves at a velocity higher than slow motor velocities. This velocity is higher for lipid cargo than rigid cargo (Fig. 3d statistical significance confirmed with students-t test, p=0.03) This is because the lipid membrane allows the motors to slide on the membrane and relieve some of its tension. Therefore, in lipid cargoes, the tension between motors with different velocities will build up at a rate slower than the rate in rigid cargoes. Since the velocities of motors are dependent on the force they experience, a lower time to build tension would mean faster velocities of motors, especially fast motors. To confirm this, we then measured the average velocity by computing a weighted sum of velocities in each of these states (Fig. 3d), weighted by the corresponding fraction of time (Fig. 3c). This reproduced the correct overall cargo velocity.

Overall, the analysis of velocity contributions from the cargo recentering mechanism and delay in strain generation mechanism reveals that it is the delay in strain generation that explains the higher velocity of lipid cargoes when carried by heterogeneous teams of motors.

### The runlength of cargo increases with an increase in the fraction of slow motors

Next, we focused on other biophysical advantages of having a mixture of slow and fast motors in a team. Experimental observations indicate that the heterogeneity in motor velocity is higher than what we would expect from random fluctuations (32), suggesting there could be some functional advantages to having high heterogeneity. This is somewhat counter-intuitive since identical teams can reduce strain generation and may therefore be more efficient. However, one might also expect to have different optimal velocity and runlength requirements for different cargoes in cells. Often, the velocities and runlengths of single motors are inter-related. Motors that step slowly often remain associated with microtubules longer (37). Previous work has shown that cargo transport by teams of motors at low ATP concentration (where motors step slowly) have a higher run length (33) for both rigid and lipid cargo.

However, the presence of all or some slower motors in the team is expected to reduce the overall cargo velocity. So we expected that cells might optimize the degree of heterogeneity in order to achieve the desired runlength while also maintaining higher cargo velocity. To explore this optimality, we ran simulations with a varying fraction of slower motors and measured the runlength (Fig. 4a) and velocity (Fig. 4b) of lipid and rigid cargoes. As expected, the runlength increases and cargo velocity decreases with an increase in the fraction of slower motors for both rigid and lipid cargo (Fig. 4a). It is to be noted that the lipid cargo runlength increases significantly more than the rigid cargo case as the fraction of slower motors increases. Thus, lipid cargo can achieve a given runlength with a lower fraction of slower motors and therefore, a higher cargo velocity. To illustrate this, consider a runlength of 3*µ*m. Lipid cargo can achieve this runlength with a fraction of about 0.25, which means a cargo velocity of about 630 nm*s*^−1^. However, a rigid cargo would need all slow motors to travel this distance, which means a cargo velocity of only 350 nm*s*^−1^.

**Figure 4.**
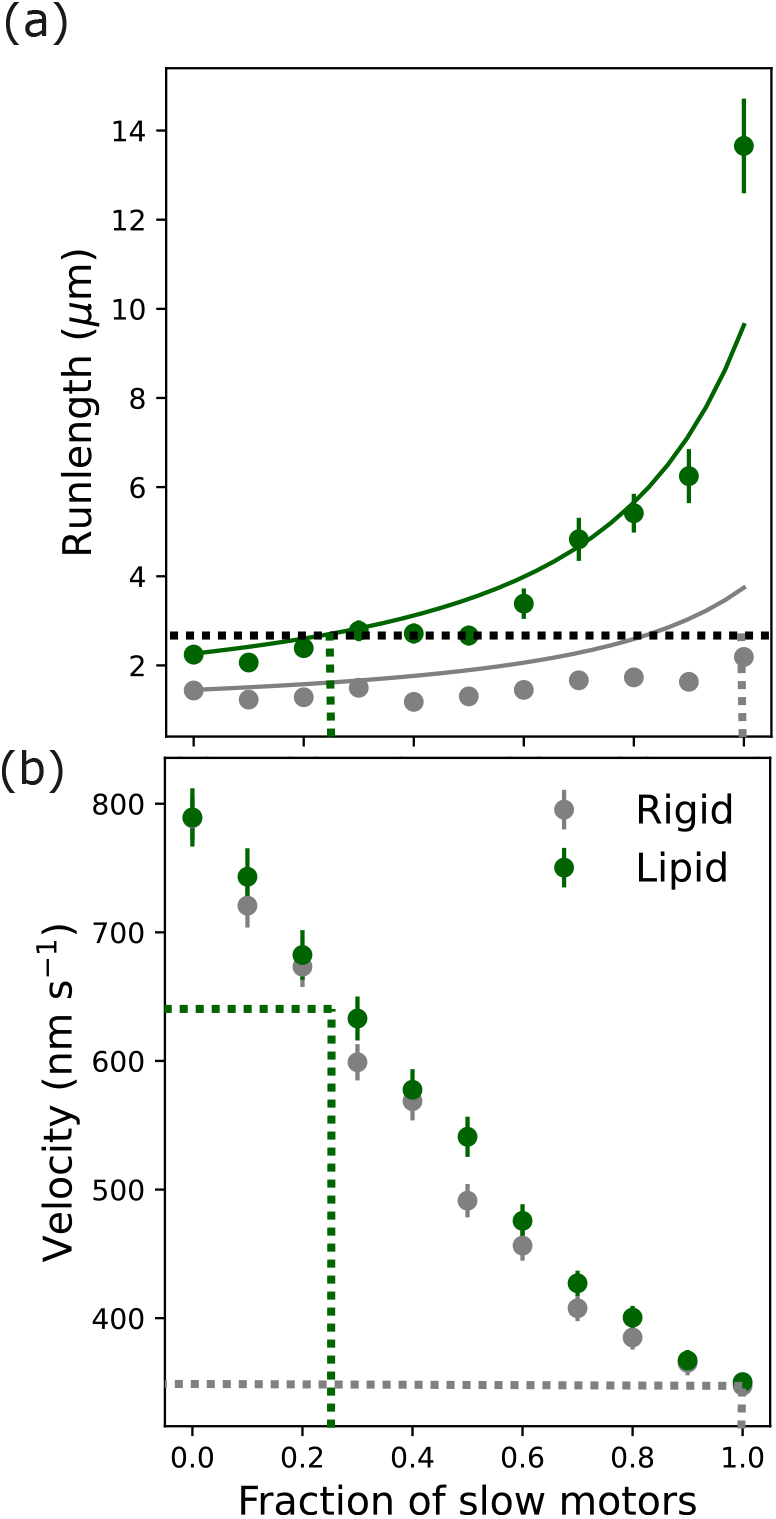
Lipid cargo achieves given runlength with a lower fraction of slower motors or with higher overall cargo velocity. (a) Runlength of the cargo. Circular markers show the mean runlength measured over 200 cargo runs. The solid lines are analytical estimations of cargo runlength by considering the weighted average of the velocity and unloaded motor unbinding rate (see supplemental information, Section III). Horizontal dashed line corresponds to a runlength of 3*µ*m and vertical dashed lines show the slow motor fractions necessary to achieve this runlength for lipid and rigid cargo. Mean velocity of cargoes. We measured velocity using the same method as we followed for the previous figures. Vertical dashed lines correspond to slow motor fractions from (a) and horizontal dashed lines show the corresponding velocities. Error bars indicate the standard error of the mean. We considered the parameters, *N* = 16 and [*AT P*]= 2 mM for these simulations same as before.

Thus, cells can tune the composition of motor teams depending on the distance that needs to be traveled and the velocity requirements. There is evidence, at least in the context of rigid cargoes, of cells making use of this tunability. These rigid cargoes are intraflagellar particles that move along axonemes in cilia. They are carried by the concerted action of kinesin-II and osm-3 kinesin motors (30, 35). In the longer middle region of cilia (about 4 *µ*m), IFT particles were observed to be carried by both slow (kinesin -II) and fast (osm-3 kinesin) motors. At the same time, the shorter distal segment (2.5 *µ*m) is found to be traversed by only fast (osm-3 kinesin) motors. Other cases where cells make use of this tunability remain to be explored.

## Conclusion

Previous experimental observations (22, 26) showed that membrane-bound cargoes (or lipid cargoes) have higher velocity than membrane-free cargoes when carried by teams of molecular motors. Even though a couple of mechanisms were proposed to explain this higher velocity of lipid cargoes, a clear quantitative explanation of the mechanisms was lacking. In this study, we show using a computational model that underlying heterogeneities in single motor velocities are crucial to observe higher velocities of lipid cargoes.Our simulations show that when motors attached to the same cargo have different velocities, they develop tension between them as they walk along the microtubules. If the motors are coupled via a membrane, the rate of this strain buildup is smaller as the attachment points are able to slide. During the time in which slow and fast motors build up tension, the cargo can move at a velocity higher than the limiting slow motor velocity. Since this time is longer in lipid cargo, the overall cargo velocity is also higher.

While velocity is one quantity that is of interest to cargo transport in cells, run length is another important quantity. Having some or all slow motors in the team enhances cargo run lengths at the expense of velocity. We find that the runlnegth of lipid cargo increases more than the rigid cargo with an increase in the fraction of slow motors. Thus lipid cargoes can traverse a given distance with a lower fraction of slower motors or higher overall cargo velocity. We also estimated the runlnegth of cargoes with a modified Klumpp expression (15, 33) (see supplemental information, Section III). This analytical model seemed to adequately predict the runlengths of rigid and lipid cargoes allowing us to extend the predictions to various other parameter regimes in future studies. Overall, our work elucidates the emergence of cooperativity between heterogeneous teams of motors in cells because of purely physical mechanisms arising from how they are coupled together. We also illustrate that the degree of heterogeneity can be used as a tunability parameter to achieve desired runlength and velocity. These predictions need to be verified with experiments of membrane-bound cargo transport by teams of motors with varying levels of heterogeneity.

Our model makes several assumptions, some of which may impact the behavior of the cargo-motor system. We assumed a simple mechanistic motor binding event followed by a diffusive search, but recent studies highlight the importance of considering the motors’ chemical state also in these bindings (27). Another crucial assumption in our model is that the detachment rates of motors decrease with a decrease in motor velocities. We assumed that all the motors are kinesin-I motors, and the rate of the detachment of motors from microtubules is expected to be directly proportional to the velocity of motors. However, cellular cargoes can have motor teams that are made up of different motor types, with different relations between runlengths and velocities. It remains to be seen what effects such heterogeneities can have on transport by teams of motors and whether they provide any functional advantages *in vivo*.

## Author Contributions

NS and AG designed the research. NS carried out the simulations and gathered data. NS and AG analyzed the data. NS and AG wrote the article.

## Acknowledgments

This work was supported by the National Science Foundation (NSF-DMS-1616926 to AG) and NSF-CREST: Center for Cellular and Biomolecular Machines at UC Merced (NSF-HRD-1547848 and NSF-HRD-2112675 to AG). AG and NS acknowledge support from the NSF Center for Engineering Mechanobiology grant (NSF-CMMI-1548571) and computing time on the Multi-Environment Computer for Exploration and Discovery (MERCED) cluster at UC Merced (NSF-ACI-1429783). NS acknowledges Graduate Dean’s dissertation fellowship from the University of California, Merced.

## Notes

### Competing Interest Statement

The authors have declared no competing interest.

## References

1. D. Ando, N. Korabel, K. C. Huang, and A. Gopinathan. Cytoskeletal network morphology regulates intracellular transport dynamics. Biophysical journal, 109(8):1574–1582, 2015.

2. D. Ando, M. Mattson, J. Xu, and A. Gopinathan. Co-operative protofilament switching emerges from inter-motor interference in multiple-motor transport. Sci. Rep., 4:7255, 2014.

3. G. Arpag, S. Shastry, W. O. Hancock, and E. Tüzel. Transport by populations of fast and slow kinesins uncovers novel family-dependent motor characteristics important for in vivo function. Biophys. J., 107(8):1896–1904, 2014.

4. B. M. Bensel, S. Previs, C. Bookwaler, K. M. Trybus,S. Walcott, and D. M. Warshaw. Spatial relationships matter: Kinesin-1 molecular motors transport liposome cargo through 3d microtubule intersections in vitro. bioRxiv, pages 2023–12, 2023.

5. B. M. Bensel, S. B. Previs, C. Bookwalter, K. M. Trybus,S. Walcott, and D. M. Warshaw. Kinesin-1-transported liposomes prefer to go straight in 3d microtubule intersections by a mechanism shared by other molecular motors. Proceedings of the National Academy of Sciences, 121(29):e2407330121, 2024.

6. M. Bovyn, B. R. Janakaloti Narayanareddy, S. Gross, and J. Allard. Diffusion of kinesin motors on cargo can enhance binding and run lengths during intracellular transport. Mol. Biol. Cell., 32(9):984–994, 2021. PMID: 33439674.

7. J. W. Driver, A. R. Rogers, D. K. Jamison, R. K. Das, A. B. Kolomeisky, and M. R. Diehl. Coupling between motor proteins determines dynamic behaviors of motor protein assemblies. Phys. Chem. Chem. Phys., 12(35):10398, aug 2010.

8. A. I. D’Souza, R. Grover, G. A. Monzon, L. Santen, and S. Diez. Vesicles driven by dynein and kinesin exhibit directional reversals without regulators. Nature Communications, 14(1):7532, 2023.

9. M. W. Gramlich, L. Conway, W. H. Liang, J. A. Labastide, S. J. King, J. Xu, and J. L. Ross. Single molecule investigation of kinesin-1 motility using engineered microtubule defects. Scientific Reports, 7(1):1–12, 3 2017.

10. R. Grover, J. Fischer, F. W. Schwarz, W. J. Walter,P. Schwille, and S. Diez. Transport efficiency of membrane-anchored kinesin-1 motors depends on motor density and diffusivity. Proc. Natl. Acad. Sci. U.S.A., 113(46):E7185–E7193, nov 2016.

11. W. O. Hancock. Bidirectional cargo transport: moving beyond tug of war. Nature Reviews Molecular Cell Biology, 15(9):615–628, Aug. 2014.

12. J. Howard. Mechanics of motor proteins and the cytoskeleton. Sinauer Associates (OUP), 2001.

13. J. Hu, S. Jafari, Y. Han, A. J. Grodzinsky, S. Cai, and M. Guo. Size- and speed-dependent mechanical behavior in living mammalian cytoplasm. Proceedings of the National Academy of Sciences, 114(36):9529–9534, Aug. 2017.

14. R. Jiang, S. Vandal, S. Park, S. Majd, E. Tüzel, and W. O. Hancock. Microtubule binding kinetics of membrane-bound kinesin-1 predicts high motor copy numbers on intracellular cargo. Proc. Nat. Acad. Sci. U.S.A., 116(52):26564–26570, 2019.

15. S. Klumpp and R. Lipowsky. Cooperative cargo transport by several molecular motors. Proc. Natl. Acad. Sci. U.S.A., 102(48):17284–17289, 2005.

16. A. Kunwar and A. Mogilner. Robust transport by multiple motors with nonlinear force– velocity relations and stochastic load sharing. Phys. Biol., 7(1):16012, 2010.

17. A. Kunwar, S. K. Tripathy, J. Xu, M. K. Mattson, P. Anand,R. Sigua, M. Vershinin, R. J. McKenney, C. C. Yu, A. Mogilner, and S. P. Gross. Mechanical stochastic tug-of-war models cannot explain bidirectional lipid-droplet transport. Proc. Natl. Acad. Sci. U.S.A., 108(47):18960–18965, 2011.

18. A. Kunwar, M. Vershinin, J. Xu, and S. P. Gross. Stepping, Strain Gating, and an Unexpected Force-Velocity Curve for Multiple-Motor-Based Transport. Curr. Biol., 18(16):1173–1183, 8 2008.

19. J. A. Labastide, D. A. Quint, R. K. Cullen, B. Maelfeyt, J. L. Ross, and A. Gopinathan. Non-specific cargo– filament interactions slow down motor-driven transport. The European Physical Journal E, 46(12):134, 2023.

20. V. Levi, A. S. Serpinskaya, E. Gratton, and V. Gelfand. Organelle transport along microtubules in xenopus melanophores: Evidence for cooperation between multiple motors. Biophysical Journal, 90(1):318–327, 2006.

21. Q. Li, J. T. Ferrare, J. Silver, J. O. Wilson, L. Arteaga-Castaneda, W. Qiu, M. Vershinin, S. J. King, K. C. Neuman, and J. Xu. Cholesterol in the cargo membrane amplifies tau inhibition of kinesin-1-based transport. Proceedings of the National Academy of Sciences, 120(3):e2212507120, 2023.

22. Q. Li, K.-F. Tseng, S. J. King, W. Qiu, and J. Xu. A fluid membrane enhances the velocity of cargo transport by small teams of kinesin-1. J. Chem. Phys., 148(12):123318, mar 2018.

23. J. Lopes, D. A. Quint, D. E. Chapman, M. Xu, A. Gopinathan, and L. S. Hirst. Membrane mediated motor kinetics in microtubule gliding assays. Sci. Rep., 9(1), July 2019.

24. F. L. Memarian, J. D. Lopes, F. J. Schwarzendahl, M. G. Athani, N. Sarpangala, A. Gopinathan, D. A. Beller, K. Dasbiswas, and L. S. Hirst. Active nematic order and dynamic lane formation of microtubules driven by membrane-bound diffusing motors. Proceedings of the National Academy of Sciences, 118(52), 2021.

25. S. S. Mogre, A. I. Brown, and E. F. Koslover. Getting around the cell: physical transport in the intracellular world. Physical Biology, 17(6):061003, Oct. 2020.

26. S. R. Nelson, K. M. Trybus, and D. M. Warshaw. Motor coupling through lipid membranes enhances transport velocities for ensembles of myosin Va. Proc. Natl. Acad. Sci. U.S.A., 111(38):E3986–E3995, 2014.

27. T. Nguyen, B. R. J. Narayanareddy, S. P. Gross, and C. E. Miles. Competition between physical search and a weak-to-strong transition rate-limits kinesin binding times. PLOS Computational Biology, 20(5):e1012158, 2024.

28. C. M. O’Connor and J. U. Adams. Essentials of cell biology. NPG Education, Cambridge, MA, 2010.

29. G. Ou, O. E. Blacque, J. J. Snow, M. R. Leroux, and J. M. Scholey. Functional coordination of intraflagellar transport motors. Nature, 436(7050):583–587, July 2005.

30. X. Pan, G. Ou, G. Civelekoglu-Scholey, O. E. Blacque,N. F. Endres, L. Tao, A. Mogilner, M. R. Leroux, R. D. Vale, and J. M. Scholey. Mechanism of transport of IFT particles in C. elegans cilia by the concerted action of kinesin-II and OSM-3 motors . Journal of Cell Biology, 174(7):1035–1045, 09 2006.

31. A. Rai, D. Pathak, S. Thakur, S. Singh, A. K. Dubey, and R. Mallik. Dynein clusters into lipid microdomains on phagosomes to drive rapid transport toward lysosomes. Cell, 164(4):722–734, Feb. 2016.

32. B. J. Reddy, S. Tripathy, M. Vershinin, M. E. Tanenbaum,J. Xu, M. Mattson-Hoss, K. Arabi, D. Chapman, T. Doolin,C. Hyeon, and S. P. Gross. Heterogeneity in kinesin function. Traffic, 18(10):658–671, 2017.

33. N. Sarpangala and A. Gopinathan. Cargo surface fluidity can reduce inter-motor mechanical interference, promote load-sharing and enhance processivity in teams of molecular motors. PLOS Computational Biology, 18(6):1–32, 06 2022.

34. N. Sarpangala, B. Randell, A. Gopinathan, and O. Kogan. Tunable intracellular transport on converging microtubule morphologies. Biophysical Reports, 4(3), 2024.

35. J. J. Snow, G. Ou, A. L. Gunnarson, M. R. S. Walker, H. M. Zhou, I. Brust-Mascher, and J. M. Scholey. Two anterograde intraflagellar transport motors cooperate to build sensory cilia on c. elegans neurons. Nature Cell Biology, 6(11):1109–1113, Oct. 2004.

36. J. O. Wilson, A. D. Zaragoza, and J. Xu. Tuning ensemble-averaged cargo run length via fractional change in mean kinesin number. Physical biology, 18(4):046004, 2021.

37. J. Xu, Z. Shu, S. J. King, and S. P. Gross. Tuning Multiple Motor Travel via Single Motor Velocity. Traffic, 13(9):1198–1205, 2012.

38. S. Yadav and A. Kunwar. Sliding of motor tails on cargo surface due to drift and diffusion affects their team arrangement and collective transport. Physical Biology, 20(1):016002, 2022.

