## Supplemental Information for "Membrane-bound cargo carried by teams of motors with heterogeneous velocities go faster and further than rigid cargo"

#### Contents

**I Brownian dynamics model** 1

**II Estimation of cargo velocity assuming single motor transport** 4

**III Analytical estimation of runlength** 4

**IV Supplemental figures** 5

#### I Brownian dynamics model

The Brownian dynamics model used in this work is similar to the model we used in our previous work (31), the only difference being that in the current model, we assign different velocities to motors. Our Brownian dynamics model builds upon various previous works (4, 7–9, 13, 17, 20, 23–26, 28–30, 35, 39). Our model is very similar to the work in Bovyn et. al. (9).

In our model, we consider a spherical cargo of radius, $R$ ($R$ = 250 nm in most cases) which has a given number of molecular motors ($N$) distributed on the surface. $N$ doesn't change during the cargo run, meaning there is no binding and unbinding of motors between the cargo surface and the solution. Each of these $N$ motors is initially assigned a random, uniformly distributed anchor position on the cargo surface. Molecular motors on rigid cargo are fixed on the cargo surface. So a given initial configuration of motors on the rigid cargo surface persists throughout the cargo run whereas motors on lipid cargo diffuse on the surface.

All $N$ motors are kinesin motors which have a rest length of $L_{mot}$ = 57 nm (27). We assume that an unbound kinesin motor binds to the microtubule with a constant rate, $\pi_0$ = 5 $s^{-1}$ (26), if some part of the microtubule is within $L_{mot}$ distance from the anchor position of that unbound motor. In other words, unbound motors bind at a specific rate to the microtubule if they can access the microtubule. The number of motors on the cargo, ($N$) is assumed to be constant, similar to several other modeling studies (9, 11). In other words, we assume the timescale for motor detachment from the cargo is much larger than the lifetime of the cargo on the microtubule.

Each cargo run is initiated with at least one motor bound to microtubule and stopped when all the motors detach from the microtubule.

A microtubule bound motor is characterized by two position vectors, anchor point ($\vec{A}$) on the cargo surface and head ($\vec{H}$) on the microtubule, and an unloaded velocity $v_{o,mot}$. To a given motor, the unloaded velocity $v_{o,mot}$ is assigned during the start of the cargo run and remains constant for the rest of the run. The head position is defined just by its continuous position along the x-axis, i.e. we do not model the lattice structure of the microtubule. We justify this based on the fact that the microtubule has multiple protofilaments and when two kinesins are on different protofilaments they can have same x-position on the microtubule. There are also models that have explicitly incorporated the fact the motor heads cannot step on each other (12, 28), which is likely of more importance when the filament has only a few tracks like actin, which has only two protofilaments in a helix, compared to 10-15 for microtubules.

A microtubule bound motor is assumed to exert a spring-like force when the length of motor exceeds the motor's rest length ($L_{mot}$) with a force constant, $k_{mot}$ = 0.32 pN/nm (14, 15). Let $\vec{L} = \vec{A} - \vec{H}$ and $L = |\vec{A} - \vec{H}|$. The force exerted by the motor on the cargo is then given by,

$$\vec{F} = -k_{mot}(L - L_{mot})\hat{L} \quad L > L_{mot} \tag{1}$$

$$= 0 \quad L \leq L_{mot} \tag{2}$$

The translation velocity of the center of mass of cargo is given by the (overdamped) Langevin equation

$$\frac{d\vec{X}}{dt} = \frac{1}{\gamma_c}\left[\sum_{j=1}^{N} \vec{F}_j + \vec{F}_{steric}\right] + \vec{\zeta} \tag{3}$$

Here, $\vec{F}_j$ is the force exerted on cargo by the j$^{th}$ motor. $\vec{F}_{steric}$ is the spring-like steric force on the cargo from the microtubule, represented with a high force constant $10k_{mot}$. We note that if the vesicle is deformable, the steric spring constant could be significantly smaller. This force is present only if the cargo-microtubule distance is less than the sum of the cargo and microtubule radii. $\vec{\zeta}$ is the random force on the cargo due to collisions with the intracellular medium. We assume a normally distributed noise with zero mean, $\langle\vec{\zeta}\rangle = 0$ and the fluctuation-dissipation relation, $\langle\zeta_\mu(t)\zeta_\nu(t')\rangle = 2\gamma_c k_B T\delta_{\mu\nu}\delta(t-t')$. $\gamma_c$ is the friction co-efficient for the cargo given by $\gamma_c = 18\pi\eta_v R$, where $\eta_v$ is the co-efficient of viscosity of cytoplasm experienced by cargo. We approximated $\eta_v$ to be equal to the co-efficient of viscosity of water in room temperature ( $20^0C$), $\eta_v = 10^{-3}$ Pa.s. We integrate Eq. 3 using the Euler-Maruyama scheme

$$\vec{X}(t+\Delta t) = \vec{X}(t) + \frac{\Delta t}{\gamma_c}\left[\sum_{j=1}^{N} \vec{F}_j + \vec{F}_{steric}\right] + \sqrt{2\gamma_c k_B T\Delta t}\vec{\xi} \tag{4}$$

where $\vec{\xi} = (\xi_1, \xi_2, \xi_3)$ and $\xi_1, \xi_2, \xi_3$ are drawn from normal distribution with zero mean and unit variance.

In addition to translational motion, the cargo also has rotational dynamics due to thermal fluctuations and torques from motor forces. Consider a motor at position $\vec{A}_i$ exerting a force $\vec{F}_i$ on the cargo. The torque on the cargo due to this motor is $\vec{r}_i \times \vec{F}_i$. The total torque on the cargo due to all the motor forces is

$$\vec{\tau} = \sum_{i=1}^{N} \vec{r}_i \times \vec{F}_i \tag{5}$$

The angular displacement in time $\Delta t$ taking into account this torque and thermal fluctuations is

$$\Delta\vec{\theta} = \frac{\vec{\tau}}{\gamma_R}\Delta t + \alpha\sqrt{4D_R\Delta t}\,\hat{n} \tag{6}$$

Where, $\gamma_R$ is the friction co-efficient, given by $8\pi\eta_v R^3$. $D_R$ is the rotational diffusion constant of the cargo. $\alpha$ is a calibration constant to match the experimentally measured rotational mean square displacement. We set the rotational diffusion constant ($D_R$) for lipid cargo to be equal to that of a free spherical bead in solution, which is $k_B T/\gamma_R = k_B T/8\pi\eta_v R^3$. The rotational diffusion constant for rigid cargo bound by one motor was measured (19) to be $7\times10^{-2}$ rad$^2 s^{-1}$. In our simulations we used this value to calibrate the rotational diffusion of rigid cargo. $\hat{n}$ in Eq. 6 is given by $\hat{n} = (n_1, n_2, n_3)$ which is a random unit vector in 3-dimensions. To get this, we draw numbers a and b from uniform distribution in the interval [0,1]. Calculate angles $\theta_t = cos^{-1}(2a-1)$ and $\phi_t = 2\pi b$. Then $\hat{n} = (n_1, n_2, n_3) = (sin\,\theta_t cos\,\phi_t, sin\,\theta_t sin\,\phi_t, cos\,\theta_t)$.

Let $\Delta\theta$ be the magnitude and $\hat{\omega} = (\,\omega_x,\,\omega_y,\,\omega_z\,)$ be the direction of this angular displacement vector $\Delta\vec{\theta}$. The Rodrigues' rotation matrix corresponding to this rotation is (34)

$$R_{\hat{\omega}}(\Delta\theta) = \mathbb{I} + \tilde{\omega}\sin\Delta\theta + \tilde{\omega}^2\,(1-\cos\Delta\theta) \tag{7}$$

where $\mathbb{I}$ is the $3\times3$ identity matrix and $\tilde{\omega}$ is given by

$$\tilde{\omega} = \begin{bmatrix} 0 & -\omega_z & \omega_y \\ \omega_z & 0 & -\omega_x \\ -\omega_y & \omega_x & 0 \end{bmatrix} \tag{8}$$

We update each anchor point position using this matrix

$$\vec{A}_i = \vec{X} + R_{\hat{\omega}}(\Delta\theta)\left(A_i - \vec{X}\right) \tag{9}$$

At each time step we also update the anchor positions of each motor on the lipid cargo surface using a similar Brownian dynamics formalism given by

$$\vec{A}(t + \Delta t) = \vec{A}(t) + \Delta l_\theta \hat{\theta} + \Delta l_\phi \hat{\phi} \tag{10}$$

where $\Delta l_\theta$ and $\Delta l_\phi$ are the small displacements along $\hat{\theta}$ and $\hat{\phi}$ directions in the plane tangential to cargo surface at $\vec{A}(t)$. $\Delta l_\theta$ and $\Delta l_\phi$ are given by

$$\begin{pmatrix} \Delta l_\theta \\ \Delta l_\phi \end{pmatrix} = \sqrt{2D\Delta t} \begin{pmatrix} \xi_a \\ \xi_b \end{pmatrix} + \frac{\Delta t}{\gamma_s} \begin{pmatrix} F_\theta \\ F_\phi \end{pmatrix} \tag{11}$$

where $\xi_a$ and $\xi_b$ are random variables obtained from normal distribution with zero mean and unit variance. $D$ is the diffusion constant for motor diffusion on cargo surface and $\gamma_s$ is the friction coefficient given by $\gamma_s = k_B T/D$. $(F_\theta, F_\phi)$ are the components of motor forces along $\hat{\theta}$ and $\hat{\phi}$ respectively which can be obtained from the motor force in Cartesian co-ordinates, $\vec{F} = (F_x, F_y, F_z)$ as follows

$$\begin{pmatrix} F_\theta \\ F_\phi \end{pmatrix} = \begin{pmatrix} \cos\theta\cos\phi & \cos\theta\sin\phi & -\sin\theta \\ -\sin\phi & \cos\phi & 0 \end{pmatrix} \begin{pmatrix} F_x \\ F_y \\ F_z \end{pmatrix} \tag{12}$$

At every time step, each bound kinesin motor hydrolyses an ATP molecule with certain probability and attempts to move forward on the microtubule. This stepping probability is a function of the motor force and also the ATP concentration. We have adopted the following relation for the stepping probability (2)

$$p_{step}(\vec{F}) = 1 - e^{-v\Delta t/\delta} \tag{13}$$

where $\delta$ is the step size, the distance moved by motor after hydrolyzing one ATP molecule ($\delta$ = 8 nm (6, 16, 38)). $v$ is the velocity of kinesin motor motor force (and also ATP concentration but that is not relevant for this study).

$v(\vec{F})$ gives the force dependence of the velocity. In the hindering direction, we assume (2, 23)

$$\begin{aligned} v_{hind}(\vec{F}) &= v_{o,mot}\left[1 - \left(\frac{F}{F_s}\right)^w\right] && F < F_s \\ &= 0 && F \geq F_s \end{aligned} \tag{14}$$

$v_{o,mot}$ is the velocity of motor when it doesn't experience any external load. For a given motor, the value of $v_{o,mot}$ is drawn from single motor velocity distribution. F is the magnitude of motor force, $F = |\vec{F}|$. $F_s$ is the stall force, the value of force beyond which kinesin motor stops walking. We considered $F_s$ = 7 pN (1, 5, 10) and $w = 2$ (24). In the assistive direction, velocity is assumed to be independent of force magnitude, $v_{asst}(\vec{F}) = v_{o,mot}$ (1, 2).

Experimentally it is found that a kinesin motor is more likely to detach from the microtubule when one head is detached from the microtubule while trying to take a step than when both the heads are bound to the microtubule (33, 36, 37). In our model we assume that a motor can detach only when it tries to take a step. At every time step we first check whether a bound motor tries to make a step with probability $p_{step}([\text{ATP}], \vec{F})$ using Eq. 13. If it tries to take a step, we check whether it detaches from the microtubule before completing the step using a microscopic off-rate whose value is calibrated based on the experimentally observed off-rate as a function of force, $F$, at saturating ATP concentrations.

$$\epsilon_{micro}(\vec{F}) = \frac{\epsilon_{obs}(\vec{F})}{p_{step}([\text{ATP}]=2\,\text{mM}, \vec{F})} \tag{15}$$

$p_{step}([\text{ATP}]=2\,\text{mM}, \vec{F})$ is the probability to step forward in time step $\Delta t$ at high ATP concentration of 2 mM. Note that as per this relation, the effective detachment rate of a motor is inversely proportional to its unloaded velocity $v_{o,mot}$.

$\epsilon_{obs}(\vec{F})$ depends on the magnitude of forces and also their directions. For hindering forces, we used the following relationship between observed off-rate and magnitude of motor force $F$ developed in (3, 18, 32) based on Kramer's theory (22) and used in several studies (2, 21, 23)

$$\epsilon_{obs}^{hind}(\vec{F}) = \epsilon_0 e^{F/F_d} \tag{16}$$

$\epsilon_0$ is the off-rate under no load condition. We used $\epsilon_0 = 0.79\, s^{-1}$ (1, 2). $F_d$ is the detachment force. We approximated $F_d$ to be equal to the stall force $F_s$. For assistive forces, the relationship between observed off-rate and magnitude of force is taken to be (1, 2)

$$\epsilon_{obs}^{asst}(\vec{F}) = \epsilon_0 + 1.56 \times 10^{12} F \tag{17}$$

#### II Estimation of cargo velocity assuming single motor transport

We have a population of 800 $nm/s$ and 350 $nm/s$ motors at a ratio of $p_f : p_s$=0.7:0.3. The lifetime of a motor with 800 nm/s speed is assumed to be $\tau_f = 1/0.79 = 1.26$ s. The lifetime of the slower motor with 350 nm/s speed is then, $\tau_s = 8/(0.79X3.5) = 2.89$ s. Assuming that cargo is carried by only one engaged motor, and initial binding is determined by the population of motors, we can estimate the cargo velocity as

$$v_{est} = \frac{p_f \tau_f v_f + p_s \tau_s v_s}{p_f \tau_f + p_s \tau_s} \tag{18}$$

For the parameters considered, $v_{est}$= 577.32 nm/s

#### III Analytical estimation of runlength

We use the analytical expression for runlength derived by previous works, (21), modified for cargoes of finite size with membranes (31)

$$r = \frac{v_{o,mot}}{N_a \pi_{ad}} \left[ \left( 1 + \frac{\pi_{ad}}{\epsilon_{o,mot}} \right)^{N_a} - 1 \right] \tag{19}$$

Where $v_{o,mot}$, $N_a$, $\pi_{ad}$, $\epsilon_{o,mot}$ are the motor speed, the available number of motors for transport, the single motor binding rate, and the mean motor off-rate, respectively. Considering $p_{slow}$ is slow motor fraction, $v_{o,slow}$ and $v_{o,fast}$ are slow and fast motor velocities, $\epsilon_{o,slow}$ and $\epsilon_{o,fast}$ are mean unloaded unbinding rates for slow and fast motors, we compute the mean motor speed to be $v_{o,mot} = p_{slow} v_{o,slow} + (1 - p_{slow}) v_{o,fast}$, and the mean motor unbinding rate to be $\epsilon_{o,mot} = p_{slow} \epsilon_{o,slow} + (1 - p_{slow}) \epsilon_{o,fast}$. We varied the available number of motors depending on the fluidity of the cargo surface (Please see our previous work for more information (31)). It is interesting to note that even this crude analytical estimation captures the observed runlength from simulations satisfactorily as a function of fraction of slower motors (data is shown in the main text).

### IV Supplemental figures

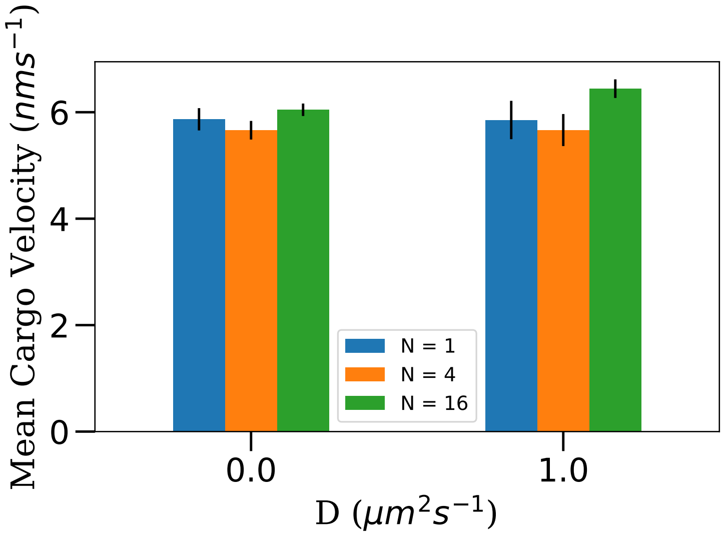


Figure S1: Velocity of cargo as a function of the number of motors on the cargo for rigid (D=0 $\mu m^2 s^{-1}$) and lipid (D=1 $\mu m^2 s^{-1}$) cargoes. Single motor velocities were 800 $nm\ s^{-1}$ and 350 $nm\ s^{-1}$ at a population ratio of 0.7:0.3. All cargo simulations were performed at saturating ATP concentration of 2 mM. 200 cargo runs were simulated in each case. For each run, the data of cargo positions were collected at a sampling rate of 100 $s^{-1}$. The cargo velocity was then calculated by measuring the mean displacement of cargo $\Delta x$ in a time interval of $\Delta t = 0.01\ s$. $v_c = \Delta x / \Delta t$

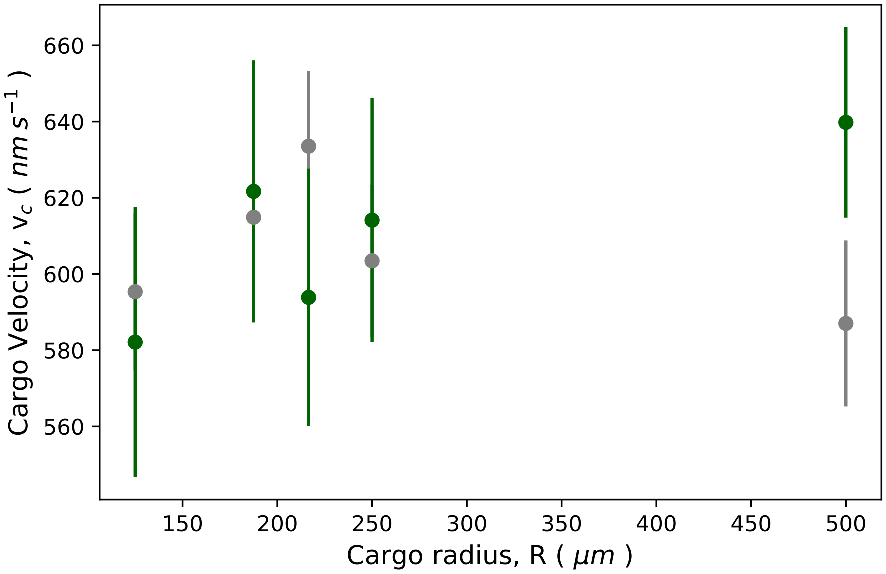


Figure S2: Velocity of cargo as a function of cargo radius for rigid (D=0 $\mu m^2 s^{-1}$, grey points) and lipid (D=1 $\mu m^2 s^{-1}$, green points) cargoes. Single motor velocities were 800 $nm\ s^{-1}$ and 350 $nm\ s^{-1}$ at a population ratio of 0.7:0.3. All cargo simulations were performed at saturating ATP concentration of 2 mM. 200 cargo runs were simulated in each case. For each run, the data of cargo positions were collected at a sampling rate of 100 $s^{-1}$. The cargo velocity was then calculated by measuring the mean displacement of cargo $\Delta x$ in a time interval of $\Delta t = 0.001\ s$. $v_c = \Delta x / \Delta t$

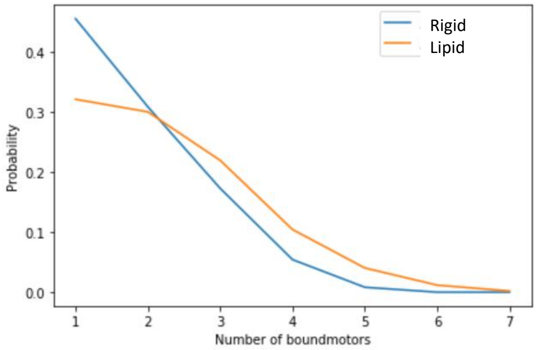


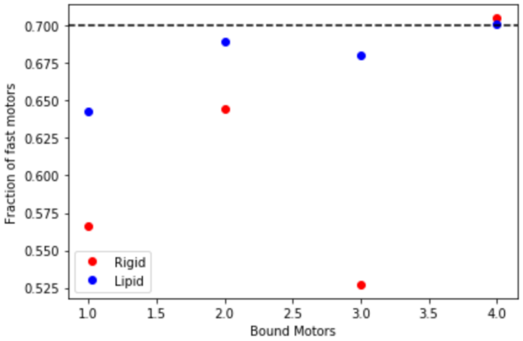


Figure S3: (Left) Probability distribution of the number of bound motors in rigid and lipid cargoes. (Right) Fraction of fast motors for a given number of bound motors. To get data in we ran simulations of cargo transport with $N = 16$ motors at [ATP]= 2 mM and recorded data at a sampling rate of 100 $s^{-1}$ for 200 cargo runs each for rigid and lipid cargoes.

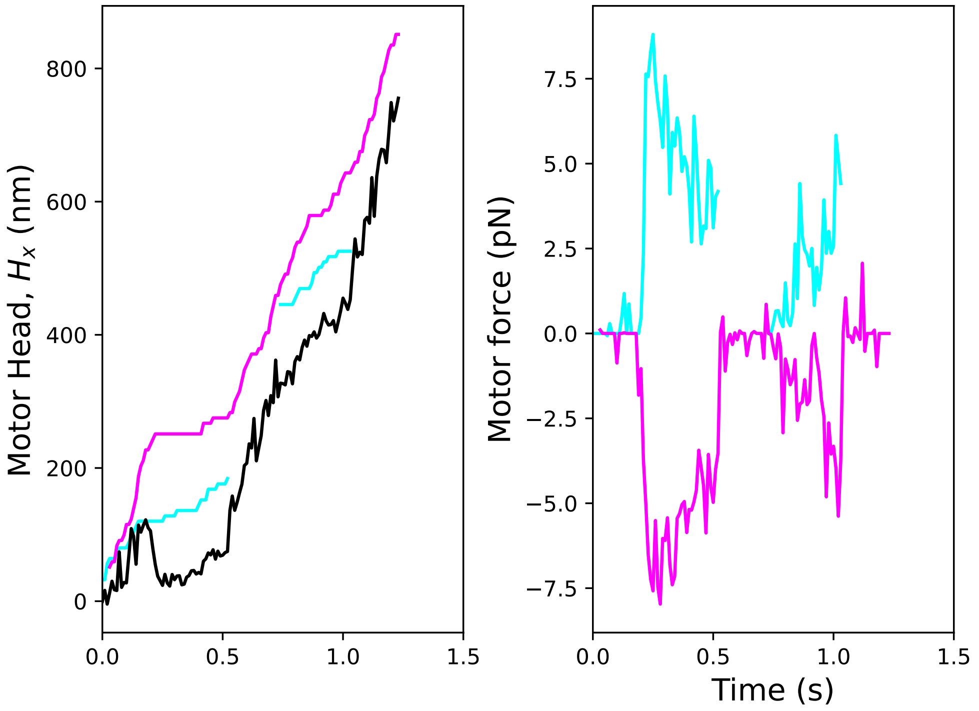


(a) Rigid Cargo ($D = 0$)

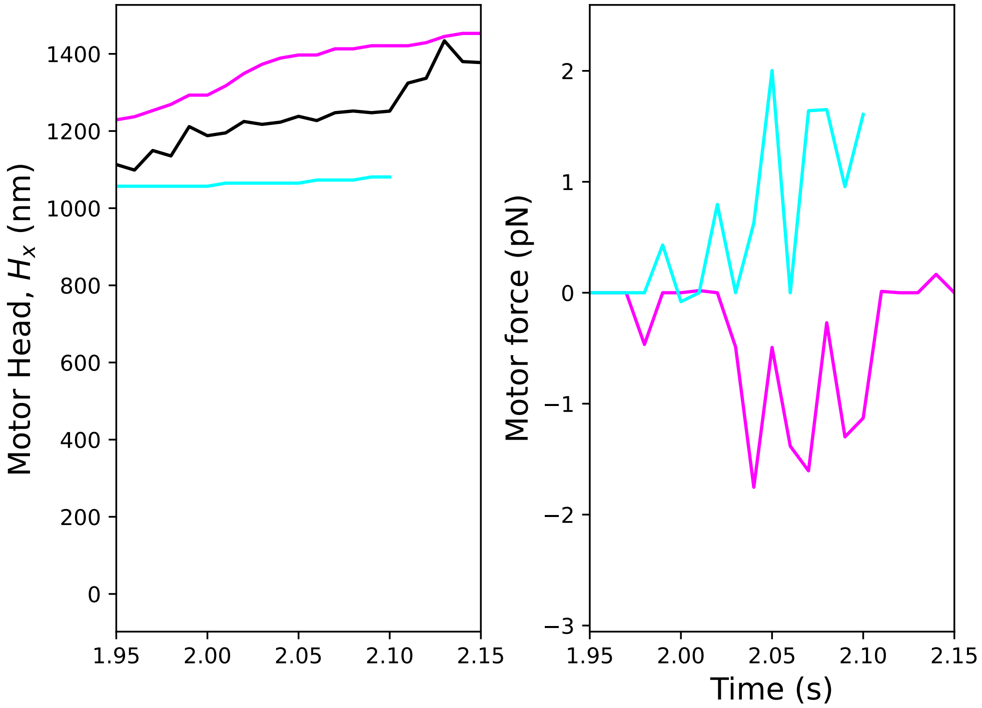


(b) Lipid Cargo ($D = 1\ \mu m^2 s^{-1}$)

Figure S4: Position and forces of motors in rigid and lipid Cargoes. Fast motors are colored in magenta; slow motors are colored in cyan. The black line represents the center of mass of the cargo. Negative force values indicate hindering forces, and positive force values indicate that the motor is experiencing assistive forces. The data were collected from cargo transport simulations with a data sampling rate of 100 $s^{-1}$. From these data, we selected a window where a fast and slow motor are simultaneously bound. The single motor velocities were 800 $nm\ s^{-1}$ (fast) and 350 $nm\ s^{-1}$ (slow) at a population ratio of 0.7:0.3.

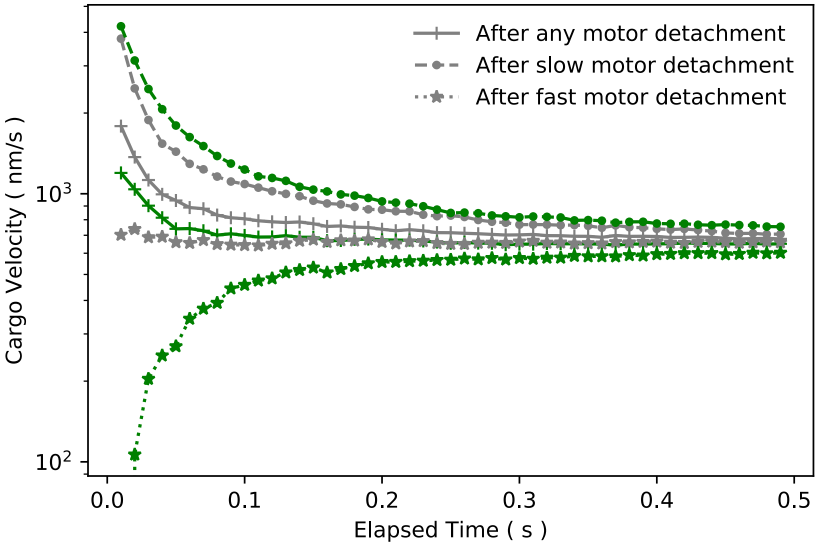


Figure S5: Average cargo velocity as a function of the time elapsed from motor detachment. We ran simulations with N=16 motors at [ATP]=2 $mM$ and recorded cargo and motor position data for 200 cargo runs each for lipid and rigid cargoes at a sampling rate of 100 $s^{1-}$. In this time series data, we identified time points where a motor detached and measured the mean displacement of cargo $\Delta x_d$, in a time window $\Delta t$ after such time points. Then we measured velocity as $v_c = \Delta x_d/\Delta t$.

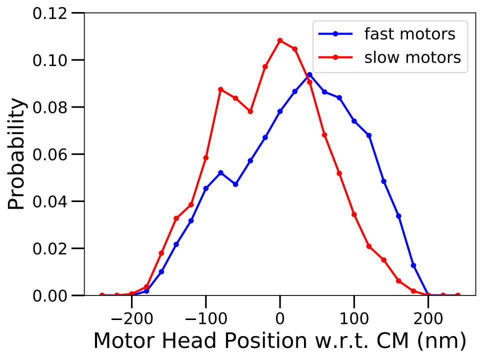


(a)

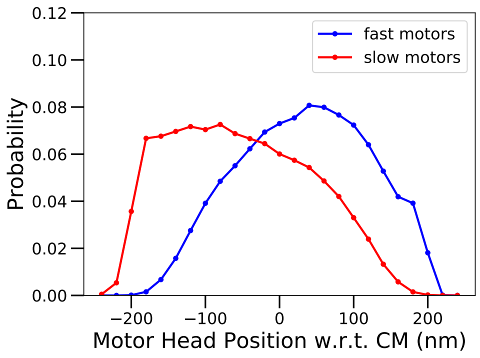


(b)

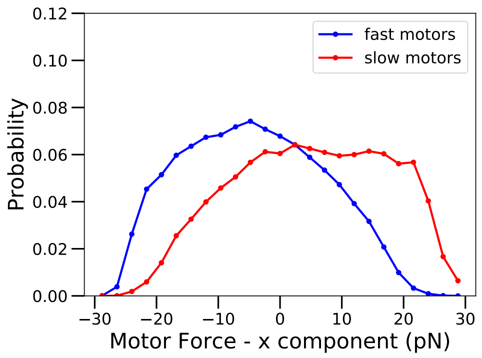


(c)

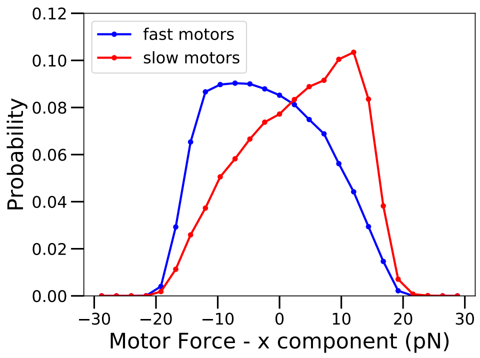


(d)

Figure S6: Distributions of motor head positions relative to the center of mass of the cargo in (a) Rigid cargo and (b) Lipid cargo. Distribution of x-component of motor force in (c) Rigid and (d) Lipid cargoes. Data obtained from cargo simulations performed at high ATP with 16 motors. Population ratio of fast (800 nm $s^{-1}$):slow (350 nm $s^{-1}$) is 0.7:0.3

### References


1. J. O. L. Andreasson. *Single-Molecule Biophysics of Kinesin Family Motor Proteins*. PhD thesis, Stanford University, Stanford,CA, 2013.

2. G. Arpag, S. Shastry, W. O. Hancock, and E. Tüzel. Transport by populations of fast and slow kinesins uncovers novel family-dependent motor characteristics important for in vivo function. *Biophys. J.*, 107(8):1896–1904, 2014.

3. G. I. Bell. Models for the specific adhesion of cells to cells. *Science*, 200(4342):618–627, May 1978.

4. J. P. Bergman, M. J. Bovyn, F. F. Doval, A. Sharma, M. V. Gudheti, S. P. Gross, J. F. Allard, and M. D. Vershinin. Cargo navigation across 3d microtubule intersections. *Proc. Natl. Acad. Sci. U.S.A.*, 115(3):537–542, 2018.

5. S. M. Block, C. L. Asbury, J. W. Shaevitz, and M. J. Lang. Probing the kinesin reaction cycle with a 2D optical force clamp. *Proc. Natl. Acad. Sci. U.S.A.*, 100(5):2351–2356, 2003.

6. S. M. Block and M. J. Schnitzer. Kinesin hydrolyses one ATP per 8-nm step. *Nature*, 388(6640):386–390, 1997.

7. S. Bouzat and F. Falo. The influence of direct motor-motor interaction in models for cargo transport by a single team of motors. *Phys. Biol.*, 7(4), 2010.

8. S. Bouzat and F. Falo. Tug of war of molecular motors: The effects of uneven load sharing. *Phys. Biol.*, 8(6), 12 2011.

9. M. Bovyn, B. R. Janakaloti Narayanareddy, S. Gross, and J. Allard. Diffusion of kinesin motors on cargo can enhance binding and run lengths during intracellular transport. *Mol. Biol. Cell.*, 32(9):984–994, 2021. PMID: 33439674.

10. N. J. Carter and R. A. Cross. Mechanics of the kinesin step. *Nature*, 435(7040):308–312, 2005.

11. K. Chen, W. Nam, and B. I. Epureanu. Effects of neighboring microtubules on the microscopic dynamics of kinesin transport. *Phys. Rev. E*, 98:052412, Nov 2018.

12. K. Chen, W. Nam, and B. I. Epureanu. Collective intracellular cargo transport by multiple kinesins on multiple microtubules. *Phys. Rev. E*, 101:052413, May 2020.

13. P. D. Chowdary, L. Kaplan, D. L. Che, and B. Cui. Dynamic clustering of dyneins on axonal endosomes: Evidence from high-speed darkfield imaging. *Biophysical Journal*, 115(2):230–241, July 2018.

14. C. M. Coppin, J. T. Finer, J. A. Spudich, and R. D. Vale. Detection of sub-8-nm movements of kinesin by high-resolution optical-trap microscopy. *Proc. Natl. Acad. Sci. U.S.A.*, 93(5):1913–1917, 1996.

15. C. M. Coppin, D. W. Pierce, L. Hsu, and R. D. Vale. The load dependence of kinesin's mechanical cycle. *Proc. Natl. Acad. Sci. U.S.A.*, 94(16):8539–8544, 1997.

16. D. L. Coy, M. Wagenbach, and J. Howard. Kinesin takes one 8-nm step for each ATP that it hydrolyzes. *J. Biol. Chem.*, 274(6):3667–3671, 1999.

17. R. P. Erickson, Z. Jia, S. P. Gross, and C. C. Yu. How molecular motors are arranged on a cargo is important for vesicular transport. *PLoS Comput. Biol.*, 7(5):e1002032, 2011.

18. E. Evans and K. Ritchie. Dynamic strength of molecular adhesion bonds. *Biophysical Journal*, 72(4):1541–1555, Apr. 1997.

19. B. Gutiérrez-Medina, A. N. Fehr, and S. M. Block. Direct measurements of kinesin torsional properties reveal flexible domains and occasional stalk reversals during stepping. *Proc. Natl. Acad. Sci. U. S. A.*, 106(40):17007–17012, 2009.

20. D. K. Jamison, J. W. Driver, A. R. Rogers, P. E. Constantinou, and M. R. Diehl. Two kinesins transport cargo primarily via the action of one motor: Implications for intracellular transport. *Biophys. J.*, 99(9):2967–2977, 2010.

21. S. Klumpp and R. Lipowsky. Cooperative cargo transport by several molecular motors. *Proc. Natl. Acad. Sci. U.S.A.*, 102(48):17284–17289, 2005.

22. H. A. Kramers. Brownian motion in a field of force and the diffusion model of chemical reactions. *Physica*, 7(4):284–304, 1940.

23. A. Kunwar and A. Mogilner. Robust transport by multiple motors with nonlinear force– velocity relations and stochastic load sharing. *Phys. Biol.*, 7(1):16012, 2010.

24. A. Kunwar, S. K. Tripathy, J. Xu, M. K. Mattson, P. Anand, R. Sigua, M. Vershinin, R. J. McKenney, C. C. Yu, A. Mogilner, and S. P. Gross. Mechanical stochastic tug-of-war models cannot explain bidirectional lipid-droplet transport. *Proc. Natl. Acad. Sci. U.S.A.*, 108(47):18960–18965, 2011.

25. A. Kunwar, M. Vershinin, J. Xu, and S. P. Gross. Stepping, Strain Gating, and an Unexpected Force-Velocity Curve for Multiple-Motor-Based Transport. *Curr. Biol.*, 18(16):1173–1183, 8 2008.

26. C. Leduc, O. Campàs, K. Zeldovich, A. Roux, P. Jolimaitre, L. Bourel-Bonnet, B. Goud, J.-F. Joanny, P. Bassereau, and J. Prost. Cooperative extraction of membrane nanotubes by molecular motors. *Proc. Natl. Acad. Sci. U.S.A.*, 101(49):17096–17101, 2004.

27. Q. Li, S. J. King, A. Gopinathan, and J. Xu. Quantitative Determination of the Probability of Multiple-Motor Transport in Bead-Based Assays. *Biophys. J.*, 110(12):2720–2728, 2016.

28. A. T. Lombardo, S. R. Nelson, M. Y. Ali, G. G. Kennedy, K. M. Trybus, S. Walcott, and D. M. Warshaw. Myosin va molecular motors manoeuvre liposome cargo through suspended actin filament intersections in vitro. *Nature Communications*, 8(1), June 2017.

29. M. J. Müller, S. Klumpp, and R. Lipowsky. Bidirectional transport by molecular motors: Enhanced processivity and response to external forces. *Biophys. J.*, 98(11):2610–2618, Jun 2010.

30. S. R. Nelson, K. M. Trybus, and D. M. Warshaw. Motor coupling through lipid membranes enhances transport velocities for ensembles of myosin Va. *Proc. Natl. Acad. Sci. U.S.A.*, 111(38):E3986–E3995, 2014.

31. N. Sarpangala and A. Gopinathan. Cargo surface fluidity can reduce inter-motor mechanical interference, promote load-sharing and enhance processivity in teams of molecular motors. *PLOS Computational Biology*, 18(6):1–32, 06 2022.

32. M. J. Schnitzer, K. Visscher, and S. M. Block. Force production by single kinesin motors. *Nat. Cell. Biol.*, 2(10):718–723, 2000.

33. A. Seitz and T. Surrey. Processive movement of single kinesins on crowded microtubules visualized using quantum dots. *EMBO J.*, 25(2):267–277, 2006.

34. B. Serge. Rodrigues' rotation formula.

35. J. O. Wilson, D. A. Quint, A. Gopinathan, and J. Xu. Cargo diffusion shortens single-kinesin runs at low viscous drag. *Sci. Rep.*, 9(1):1–12, 2019.

36. J. Xu, Z. Shu, S. J. King, and S. P. Gross. Tuning Multiple Motor Travel via Single Motor Velocity. *Traffic*, 13(9):1198–1205, 2012.

37. J. Yajima, M. C. Alonso, R. A. Cross, and Y. Y. Toyoshima. Direct Long-Term Observation of Kinesin Processivity at Low Load. *Curr. Biol.*, 12(4):301–306, 2002.

38. A. Yildiz, M. Tomishige, R. D. Vale, and P. R. Selvin. Kinesin walks hand-over-hand. *Science*, 303(5658):676–678, 2004.

39. Y. Zhang. Cargo transport by several motors. *Phys. Rev. E*, 83(1):011909, 1 2011.
